# Water stress and senescence increase calcium oxalate crystals and decrease their solubility in amaranth leaves

**DOI:** 10.64898/2026.09.17.752438

**Authors:** Ivan T. Cerritos-Castro, Alberto Barrera-Pacheco, Araceli Patrón-Soberano, Hugo M. Ramírez-Tobías, Ana P. Barba de la Rosa

## Abstract

Calcium oxalate (CaOx) crystals are widespread in plants, yet their physiological roles remain debated. Although calcium sequestration is the most widely accepted function, some studies propose that CaOx crystals can serve as a carbon source, implying calcium release. These studies report decreases in CaOx crystals under water stress and other CO₂-limiting conditions, suggesting that crystals are oxidized to CO₂ to sustain the Calvin cycle. To test this hypothesis, we subjected 39-day-old *Amaranthus cruentus* plants to 10 days of water stress, followed by rewatering and a 16-day recovery period. Crystal abundance was analyzed by polarization microscopy, first in fixed leaves and then in optically cleared leaves. In control plants, CaOx crystals were present from the leaf-primordium stage and increased throughout leaf expansion, remaining abundant until late senescence. Water stress accelerated this pattern, and rewatering had no detectable effect. Optical clearing was intended to improve crystal visualization but unexpectedly dissolved crystals in some leaves. Serendipitously, we found that crystal solubility decreased with leaf age in control plants. Water stress accelerated this loss of solubility, whereas rewatering restored solubility to levels characteristic of younger leaves. Natural senescence increased CaOx crystal abundance and rendered them increasingly recalcitrant; water stress accelerated both processes. These findings challenge the carbon-source hypothesis.

## INTRODUCTION

Calcium oxalate (CaOx) is a poorly water-soluble compound (Belliveau and Griffin, 2001). Both soluble and insoluble CaOx, in form of crystals, are widely present in plants, and crystals can constitute up to 80% of plant dry weight. They are formed in a highly regulated manner inside the vacuoles of specialized cells known as crystal idioblasts (Franceschi and Nakata, 2005). In contrast to chemically synthesized amorphous CaOx, plant CaOx crystals display specific geometric morphologies, including druses, crystal sand, styloids, and raphides (Franceschi and Nakata, 2005). Numerous physiological functions have been proposed for these crystals, including mechanical defense, calcium and oxalate homeostasis, light scattering and collection, and serving as a carbon source under water-stress conditions (Franceschi and Nakata, 2005; Korth et al., 2006; Kuo-Huang et al., 2007; Tooulakou et al., 2016a). However, aside from their mechanical defense role, their physiological significance remains controversial.

Water stress induces stomatal closure to limit transpiration, but this simultaneously restricts CO₂ uptake (Tooulakou et al., 2016a). Under such conditions, it has been proposed that CaOx crystals are dissolved and oxidized by oxalate oxidase to CO₂, which is then incorporated into the Calvin cycle to sustain photosynthesis; a mechanism called “alarm photosynthesis” (Tooulakou et al., 2016b). Several studies have reported decreases in CaOx crystals under CO₂ limitation or water stress: in *Amaranthus hybridus* leaves treated with abscisic acid (which induces stomatal closure) (Tooulakou et al., 2016b); in *A. hybridus* leaves under CO₂ starvation(Tooulakou et al., 2018); in *Colobanthus quitensis* leaves under CO₂ starvation(Gómez-Espinoza et al., 2020); in *Vitis vinifera* leaves under water stress(Kolyva et al., 2023); in *Ipomoea batatas* shoots under water stress(Gouveia et al., 2020); and in *Fagopyrum esculentum* leaves under water stress(Gaberščik et al., 2020). Moreover, in a previous work we identified chloroplast-associated proteins and structures embedded in CaOx crystals in *A. cruentus* leaves, suggesting a possible link with photosynthesis (Cerritos-Castro et al., 2022). Nevertheless, calcium homeostasis remains the most widely accepted physiological function of these crystals (Franceschi and Nakata, 2005). Calcium is essential for regulating numerous biochemical processes and enters plants readily through the roots; thus, CaOx crystals provide a high-capacity mechanism for sequestering excess calcium (Franceschi and Nakata, 2005). If CaOx degradation were used as a carbon source, the concomitant release of calcium could potentially be detrimental.

Physiological responses to water stress unfold over time, while leaves simultaneously progress through natural senescence. Leaf senescence is the terminal developmental stage and promotes nutrient remobilization from older leaves to younger tissues or reproductive organs, including carbon stores such as starch (YANG et al., 2003; Zhao et al., 2022). Although water stress induces stomatal closure, it also accelerates senescence (YANG et al., 2003). Therefore, under water stress, CaOx crystals may be subject both to the proposed “alarm photosynthesis” process and to senescence-driven nutrient redistribution. To our knowledge, this combined effect has not been investigated.

In this work, we analyze the effects of water stress and senescence on CaOx crystals in *Amaranthus cruentus* leaves. We hypothesized that if plants are subjected to water stress, their CaOx crystal abundance will decrease or disappear over time.

## RESULTS

### Experimental design and leaf responses to water stress

We grew *Amaranthus cruentus* plants in an acclimatized greenhouse (Supplementary Fig. 1). To obtain a broad overview of CaOx crystal responses to water stress and senescence, we divided the plants into three groups: control, water-stressed, and rewatered. When plants reached 39 days of age, the experiment began (day 0). The control group was watered continuously. The water-stressed group received no further irrigation. The rewatered group remained unwatered until severe signs of stress appeared (day 20), after which watering was resumed continuously.

Control plants developed normally. In stressed plants, soil humidity decreased from 16% on day 0 to 2% by day 13, when visible signs of water stress appeared and progressively worsened. Rewatered plants recovered turgidity within one day (day 20 to 21). After day 21, stressed plants were discarded, while control and rewatered plants continued to grow until day 36 (75 days old), when panicles emerged (Fig. 1).

**Figure 1.**
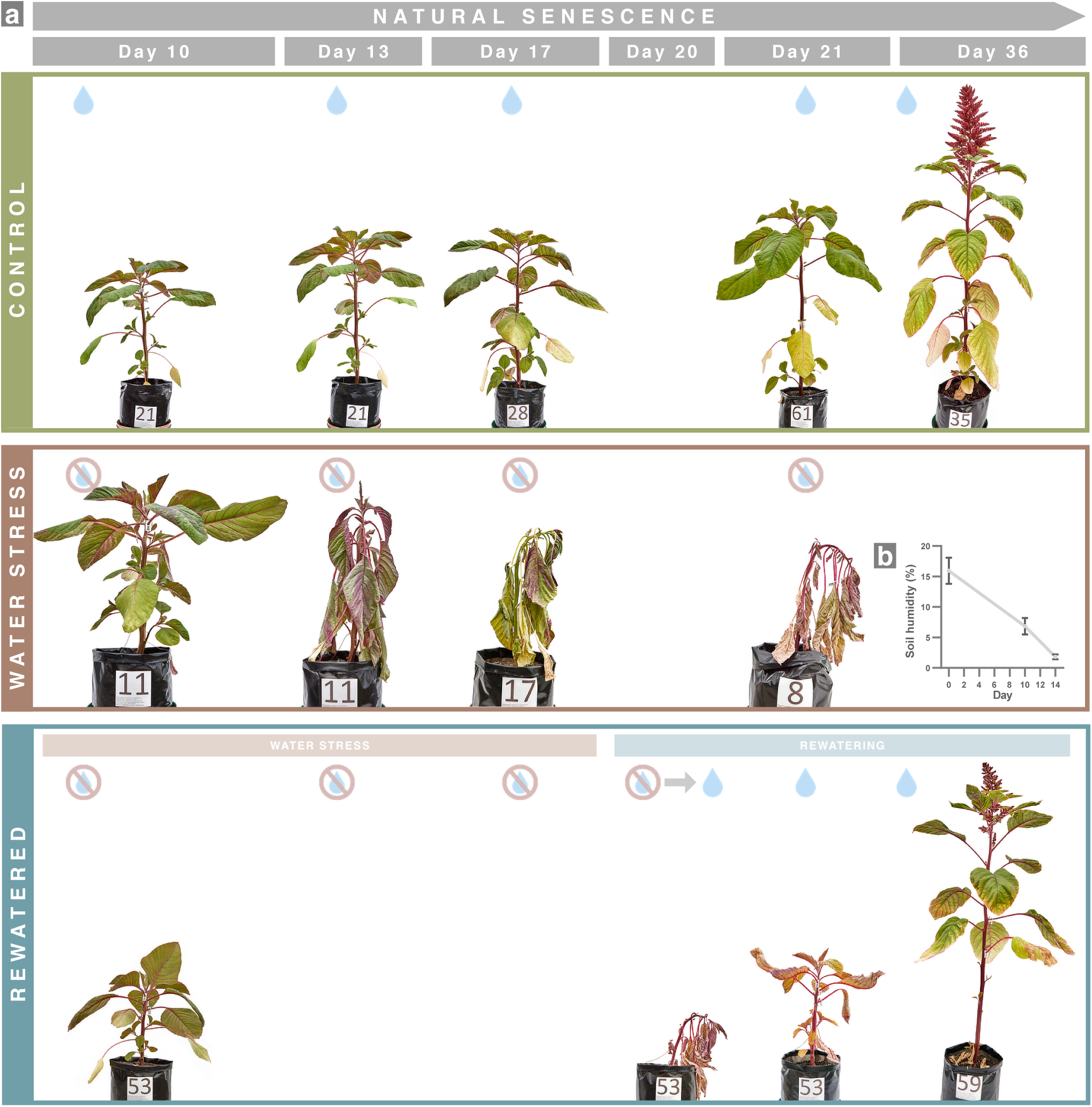
Experimental design. (a) *Amaranthus cruentus* plants at different experimental stages. (b) Soil moisture dynamics in the water-stress treatment (n = 34). 39 day-old plants (Day 0) were assigned to control, water-stress, or rewatered groups. Control plants were irrigated continuously; stressed plants received no irrigation; and rewatered plants were withheld from irrigation until severe water stress was reached (Day 20), after which irrigation resumed. Leaves were sampled on Days 13, 17, 21, and 36 (three biological replicates). Visible signs of stress appeared when soil moisture fell to ≈1–2% on Day 13 and worsened by Day 20. After rewatering, plants rapidly regained turgidity (within 24 h) and continued to grow, eventually producing panicles.

To capture the dynamic response of leaves to water stress, we analyzed the 4th, 6th, 8th, and 10th fully expanded leaves (basipetal leaf positions) throughout the experiment. These leaves were labeled on day 0 to allow accurate tracking (Fig. 2). Three biological replicates of each leaf position were sampled on days 13, 17, 21, and 36 (Supplementary Figs. 2–5). Leaf labeling enabled us to differentiate the effect of water stress from the inevitable effect of leaf aging. In this design, leaf age was determined by both leaf position and experimental day: at any given timepoint, lower leaf positions were younger, and higher leaf positions were older, but all senesced progressively across the experiment.

**Figure 2.**
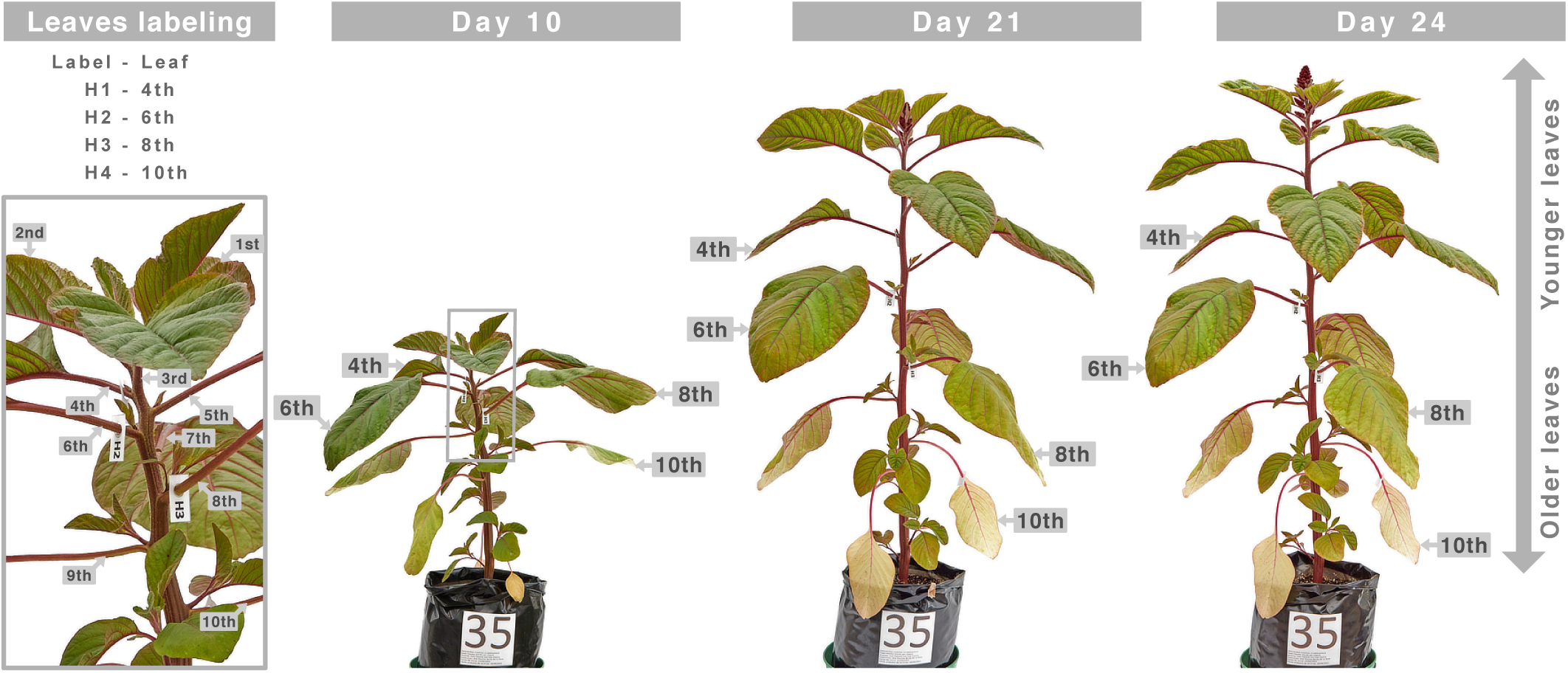
Tracking the same leaves throughout the experiment. We monitored the 4th, 6th, 8th, and 10th fully expanded leaves (basipetal leaf positions) to capture senescence and stress-related differences. Because new leaves continued to emerge, leaf position was set at Day 0 with paper tags. Older leaves (8th and 10th) senesced earlier and dried more rapidly under water stress than younger leaves (4th and 6th). Leaf age in this design is related to leaf position plus experimental day, allowing direct comparison across leaf positions and time.

Control leaves senesced gradually, with overlapping developmental stages across leaf positions. This series provided a continuum from very young, fully expanded leaves to nearly abscising chlorotic leaves. Under water stress, leaves dehydrated progressively: first the older leaves (10th and 8th), and later the younger ones (6th and 4th). By day 21, all stressed leaves exhibited severe signs such as chlorosis, wrinkling, and folding. The 4th and 6th stressed leaves remained slightly hydrated and pliable, whereas the 8th and 10th leaves became brittle. Consistent with this, only the 4th and 6th stressed leaves recovered turgidity after rewatering. Rewatered leaves showed partial recovery: tissue adjacent to major veins regained greenness, whereas distal blade regions remained chlorotic but hydrated (Supplementary Figs. 2–3). A detailed visual record is provided in Supplementary Video 1.

### Response of calcium oxalate crystals to water stress and senescence

CaOx crystals were present from the earliest stages of leaf development and continued to accumulate until late senescence. We observed abundant CaOx crystals in leaf primordia (future true leaves) of seedlings, followed by a progressive increase with leaf age in fully expanded control leaves (Supplementary Fig. 6; Fig. 3). This temporal increase was more evident in older leaves (10th) than in younger ones (4th).

**Figure 3.**
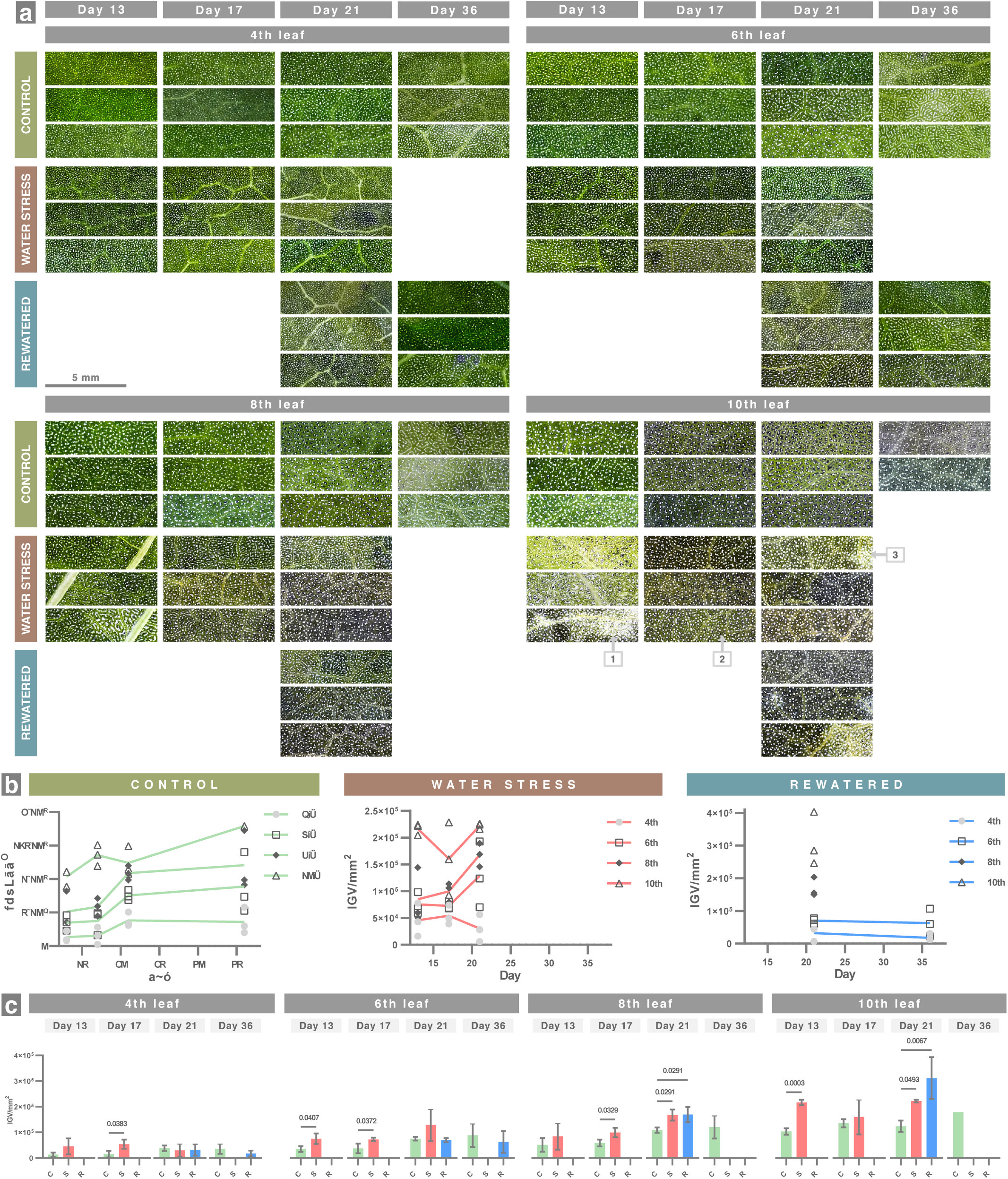
Calcium oxalate crystal abundance increased with water stress and senescence. (a) Polarized-light micrographs of fixed leaf samples at each experimental stage (triplicates shown). (b) Temporal changes in CaOx crystal abundance for each leaf position.(c) Comparison of CaOx crystal abundance among treatments at each stage. Student’s t-test (two groups) or one-way ANOVA with Tukey’s test (three groups) was used (n = 3). p < 0.05 is shown. IGV, integrated gray value; C, control; S, stress; R, rewatered. Amaranth accumulates CaOx crystals as crystal sand in veins and druses in the leaf blade. Quantification focused on the blade because veins produced excessive brightness, although both tissues showed similar treatment responses (Supplementary Data 1–2). Control leaves showed a strong positive correlation between CaOx abundance and leaf age (leaf position and experiment day). The stress treatment produced a similar but accelerated increase in the 6th and 8th leaves. However, the 4th and 10th leaves exhibited distinct behaviors. The 4th leaves showed a slight decrease over time. In contrast, the 10th leaves showed an abnormal crystal accumulation on day 13 (1), which disappeared by day 17 (2) and reappeared slightly by day 21 (3), consistent with a transient dissolution event. Rewatered leaves generally maintained the age-dependent pattern and showed little change over time. Overall, CaOx crystals increased with natural aging, were accelerated by water stress in older leaves, and may be delayed in younger leaves.

From leaf primordia to senescent leaves, druse idioblasts occurred exclusively within the intermediate mesophyll, approximately halfway through the leaf thickness. Younger leaves (day 13) contained uniformly sized druses (≈50 µm), whereas older leaves (day 36) also contained numerous smaller druses (≈10–30 µm). These smaller druses appeared during senescence and accounted for much of the increased crystal abundance (Supplementary Fig. 7).

Water stress accelerated crystal accumulation, particularly in older leaves. Stressed leaves contained significantly more crystals than their corresponding controls (Fig. 3; Supplementary Data 1). As in control plants, the 6th and 8th stressed leaves showed a progressive increase in crystal abundance over time. However, the 4th and 10th leaves exhibited distinct behaviors. The 4th leaves showed a slight decrease over time. In contrast, the 10th leaves displayed an abnormal extracellular crystal accumulation on day 13, which disappeared by day 17 and reappeared slightly by day 21 (Supplementary Fig. 8 and Supplementary Data 2).

Rewatering had no detectable effect on crystal accumulation. On day 21, one day after rewatering, rewatered leaves contained similar amounts of crystals as their corresponding stressed leaves. No changes were observed by day 36.

### Response of calcium oxalate crystals to optical clearing

Optical clearing was intended to improve crystal visualization but unexpectedly dissolved crystals in some samples, revealing solubility differences. CaOx crystal solubility decreased with both leaf age and water stress, whereas rewatering partially restored solubility to levels characteristic of younger stages.

In control plants, crystal solubility decreased with age: 10th leaves had more resistant crystals than 4th leaves, and solubility decreased progressively over time in all leaf positions. Water stress accelerated this loss of solubility in the 6th, 8th, and 10th leaves but halted it in the 4th leaves. Rewatering had no detectable effect on the 4th, 8th, and 10th leaves compared with their stressed counterparts. However, the 6th rewatered leaves on days 21 and 36 had solubility levels comparable to the 6th control leaves on day 13. In other words, crystals in the 6th leaf recovered the solubility profile of a younger pre-stress stage (Fig. 4; Supplementary Data 2).

**Figure 4.**
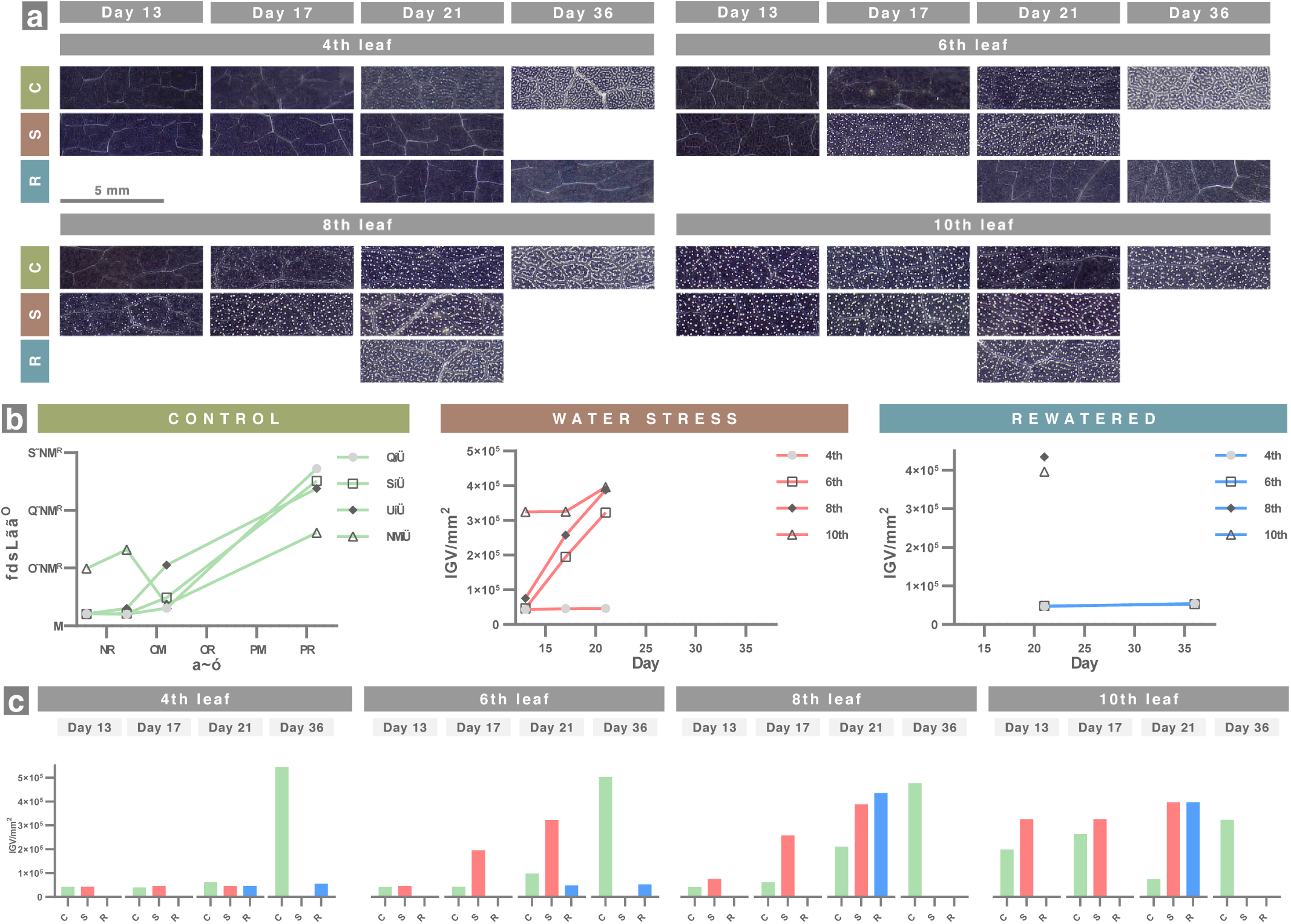
Calcium oxalate crystal solubility decreased with water stress and senescence. (a) Polarized-light micrographs of optically cleared leaf samples at each stage (one per biological replicate). (b) Temporal changes in non-dissolved CaOx crystals for each leaf position. (c) Treatment comparison at each stage. IGV, integrated gray value; C, control; S, stress; R, rewatered. Optical clearing was intended to improve visualization, but it partially dissolved CaOx crystals in some samples. Serendipitously, remaining (non-dissolved) crystals correlated positively with leaf age in the control group. Under water stress, the 6th, 8th, and 10th leaves retained up to 3.5-fold more crystals than controls, whereas the 4th leaf resembled its control counterpart. In contrast, rewatered 4th and 6th leaves retained up to 8.5-fold fewer crystals than controls. These results indicate that CaOx crystals become increasingly insoluble with aging, water stress accelerates this process in older leaves, and rewatering restores solubility in young leaves to a younger physiological state.

## DISCUSSION

Our hypothesis was incorrect: calcium oxalate (CaOx) crystals did not decrease under water stress in amaranth leaves. Instead, they increased with both water deficit and time, challenging previous reports proposing that CaOx crystals are mobilized as a carbon source under water stress conditions (Tooulakou et al., 2018, 2016b, 2016a). Rather than being consumed, our evidence shows that CaOx crystals gradually accumulate and become increasingly permanent from embryonic stages to senescence.

In control plants, CaOx crystals were already present in leaf primordia and continued increasing throughout leaf development up to senescence (Supplementary Fig. 6, Fig. 3, Fig. 4). This trend contrasts with a previous study reporting maximal crystal density at younger developmental stages (Giannopoulos et al., 2019). The tertiary veins form a mesh in which the interstices are filled by intermediate mesophyll cells and druse idioblasts. Until day 21, mostly large druses (≈50 µm) were uniformly distributed among intermediate mesophyll cells. By day 36, numerous small druses (≈10–30 µm) appeared, forming nearly continuous linear arrangements with the larger druses (Supplementary Fig. 7). We refer to these as “early druses” and “late druses,” respectively. Early druses were associated with young and early-senescent stages, whereas late druses appeared only during late senescence in all leaves. Notably, senescence progression differed by leaf position: on day 36 the 4th leaves were just beginning to yellow, whereas the 10th leaves were fully senescent and about to abscise. Thus, the emergence of late druses is not independently regulated by each leaf but reflects systemic developmental cues.

Leaf mesophyll is traditionally divided into palisade and spongy tissues, yet the cells located in the plane of the vascular bundles (central half of the blade) behaved differently. We designate these cells as intermediate mesophyll. Early druses were found exclusively among intermediate mesophyll in leaf primordia (Supplementary Fig. 6), and late druses originated from the dedifferentiation of these same cells (Supplementary Fig. 7). Druses were never observed among palisade or spongy mesophyll. Therefore, although intermediate mesophyll may resemble classic mesophyll types morphologically, it is phenotypically distinct. A similar intermediate-mesophyll behavior has been described in *Glycine max*, where it also differs physiologically from palisade and spongy mesophyll (Cerritos-Castro et al., 2025).

Water stress accelerated both CaOx accumulation and the appearance of late druses. This effect was stronger in older leaves: the 10th leaves showed the greatest increase over time relative to the 4th and 6th leaves (Fig. 3). Because older leaves dried first (Supplementary Fig. 2–5), this pattern may relate to differential desiccation dynamics. Water stress commonly induces premature senescence (YANG et al., 2003). In control leaves, CaOx accumulation naturally increased during late senescence; therefore, stress-induced CaOx accumulation may simply reflect premature activation of senescence-related pathways.

However, we observed a striking exception in the 10th stressed leaves: a transient and unusually high extracellular crystal accumulation occurred on day 13, followed by a sharp decline by day 17 (Fig. 3; Supplementary Data 1). These extracellular crystals differed from intracellular druses, which showed no such reduction (Supplementary Fig. 8). Although extracellular CaOx crystals have been reported in plants, they are uncommon (Franceschi and Nakata, 2005). Their transient nature raises mechanistic questions: How could a dehydrating leaf dissolve these crystals? Are they composed strictly of CaOx? Do they form via cellular or purely chemical processes (as opposed to the tightly regulated intracellular druses) (Cerritos-Castro et al., 2022)?

Oxalate may function as an osmotic regulator in plants (Gouveia et al., 2020). The abnormal extracellular accumulation could reflect rapid increases in soluble oxalate and calcium that drove simple chemical precipitation (Belliveau and Griffin, 2001). Intracellular druses incorporate organic matrix components during development (Cerritos-Castro et al., 2022), which likely contributes to their lower solubility. Extracellular precipitates lacking such matrix components may redissolve more readily due to chemical equilibrium shifts. This difference in organic content may be a key determinant of crystal solubility.

Rewatering did not restore turgidity in the 8th and 10th leaves, and these leaves progressed to full senescence. In contrast, 4th and 6th leaves recovered turgidity, and although differences were not statistically significant, rewatered leaves tended to show equal or fewer crystals relative to their respective control and stressed leaves (Fig. 3; Supplementary Fig. 2–5). This suggests that rewatering may reset CaOx levels to those of earlier developmental stages.

Optical clearing partially dissolved CaOx crystals despite their low solubility (Belliveau and Griffin, 2001), revealing important physiological patterns. First, CaOx solubility decreased with both age and water stress. Second, water stress halted this insolubilization process in the 4th leaf. Third, rewatering restored solubility to pre-stress levels in the 6th leaf. These findings indicate that, after formation, CaOx crystals continue to “mature”—becoming less soluble—as leaves age. Water stress can accelerate or interrupt this maturation depending on leaf position, and recovery can partially reverse it. The 8th and 10th stressed leaves, which lost turgidity and senesced (Supplementary Fig. 4–5), showed increased crystal load (Fig. 3) and reduced solubility (Fig. 4). Conversely, the recovering 6th leaves regained turgidity and greenness (Supplementary Fig. 3), displayed crystal amounts similar to controls, and restored solubility to a younger state (Fig. 3–4). Notably, in rewatered 6th leaves, some regions showed complete dissolution of CaOx whereas neighboring “dead zones” retained crystals, indicating localized physiological rejuvenation (Supplementary Data 2).

Despite decades of study, CaOx crystals remain enigmatic. Their most cited roles—calcium regulation and carbon source—are still under debate. A knockout study in *Medicago truncatula* (a natural crystal-forming species) showed that plants lacking CaOx crystals grow normally, suggesting that CaOx formation is not essential for normal development (Nakata and McConn, 2000)). Our results strongly challenge their proposed function as a water stress-mobilizable carbon reservoir. Under both natural aging and water stress, CaOx crystals increase and become more recalcitrant, which is incompatible with a role as readily available carbon stores.

Finally, our study highlights an experimental consideration often overlooked: many plant studies sample only the 4th fully expanded leaf. By using leaves of multiple developmental stages, we uncovered leaf-number-dependent differences in CaOx dynamics that would otherwise remain hidden. Future research on CaOx crystals should carefully control for leaf position and plant age, as both strongly influence crystal abundance, type, and solubility.

## METHODS

### Plant material

Seeds of *Amaranthus cruentus* L. cv. Amaranteca were germinated on BM2 substrate (Berger, Saint-Modeste, Quebec, Canada) for 20 days in an acclimatized chamber under a 12/12 h light–dark photoperiod at 25 °C. Seedlings were transplanted into 2 kg of BM2 substrate, transferred to an acclimatized greenhouse, and maintained at 15–35 °C. Temperature, relative humidity, and light intensity were monitored throughout the experiment. Soil moisture was continuously measured, and plants were irrigated, photographed, and fertilized with 1.5 g L⁻¹ vegetable nutrient solution (Hydro-Environment, Mexico).

At 39 days of age (Day 0), plants were assigned to control, water-stress, or rewatered groups. The basipetal 4th, 6th, 8th, and 10th fully expanded leaves were identified and labeled with paper tags. Control plants were continuously watered; stressed plants received no irrigation; and rewatered plants were withheld from irrigation until Day 20, when watering was resumed. To ensure equal nutrient conditions, control plants were not fertilized between Days 0 and 21. Leaf samples were collected from the middle lamina of each leaf position and treatment on Days 13, 17, 21, and 36 from three biological replicates.

### Sample analysis

Leaf samples were fixed overnight at 4 °C in 3% glutaraldehyde and 2% formaldehyde in PBS. Samples were rinsed twice in PBS at 4 °C and stored overnight in 30% ethanol. Whole-sample polarization micrographs were acquired prior to clearing (Supplementary Data 1).

One sample per biological replicate was optically cleared following(Warner et al., 2014) with modifications. Samples were incubated in 10% KOH for 21 h, followed by 39 days in clearing solution (6 M urea, 30% glycerol, 0.1% Triton X-100) on a rocking platform. Cleared samples were imaged again under polarized light (Supplementary Data 2).

For histological analysis, 5 × 5 mm subsamples were excised from the 30% ethanol-stored samples. Subsamples were processed as described previously (Cerritos-Castro et al., 2025). Briefly, samples were dehydrated through an ethanol series (50%, 70%, 90%, and 100%; 1 h each), pre-embedded in a 1:1 mixture of absolute ethanol:LR White resin overnight on a rocker, and then embedded in pure LR White resin overnight. Samples were transferred to gelatin capsules with fresh resin and polymerized at 50 °C for 48 h. Semi-thin sections (500 nm) were obtained with an ultramicrotome (RMC, Boeckeler Instruments, Tucson, AZ, USA), stained with safranin and fast green, and imaged using DIC microscopy (Zeiss Imager M2, Carl Zeiss, Oberkochen, Germany).

### Image analysis

Polarization micrographs were imported into Capture One 20, and the white balance of each image was adjusted based on calcium oxalate crystal borders. Images were converted to grayscale by setting color sensitivity to zero and further adjusted using the “luma curve” to achieve an almost completely black background. Processed images were exported to Adobe Photoshop, where a representative 4 × 4 mm region per sample was selected. Calcium oxalate crystal density was quantified using the built-in image analysis tools. Large veins were excluded due to high brightness that produced noise. Statistical analyses were performed in GraphPad Prism 9.

## Supplementary information

Supplementary video 1: https://youtu.be/isfqsHKfBd4

Supplementary Data: https://doi.org/10.5281/zenodo.22818965

## Acknowledgments

Ivan Takeshi Cerritos Castro thanks to CONACYT fellowship 719217. The authors thank to Laboratory for Nanoscience and Nanotechnology Research, LINAN, IPICYT.

## Author contributions

**Ivan Takeshi Cerritos-Castro**: Conceptualization, Methodology, Investigation, Formal analysis, Visualization, Writing – original draft. **Olga Araceli Patrón-Soberano**: Investigation, Methodology. **Alberto Barrera-Pacheco**: Investigation, Methodology. **Hugo M. Ramírez-Tobías**: Investigation, Methodology. **Ana Paulina Barba de la Rosa**: Conceptualization, Resources, Supervision, Writing – review & editing.

---

**SUPPLEMENTARY FIGURE 1.**
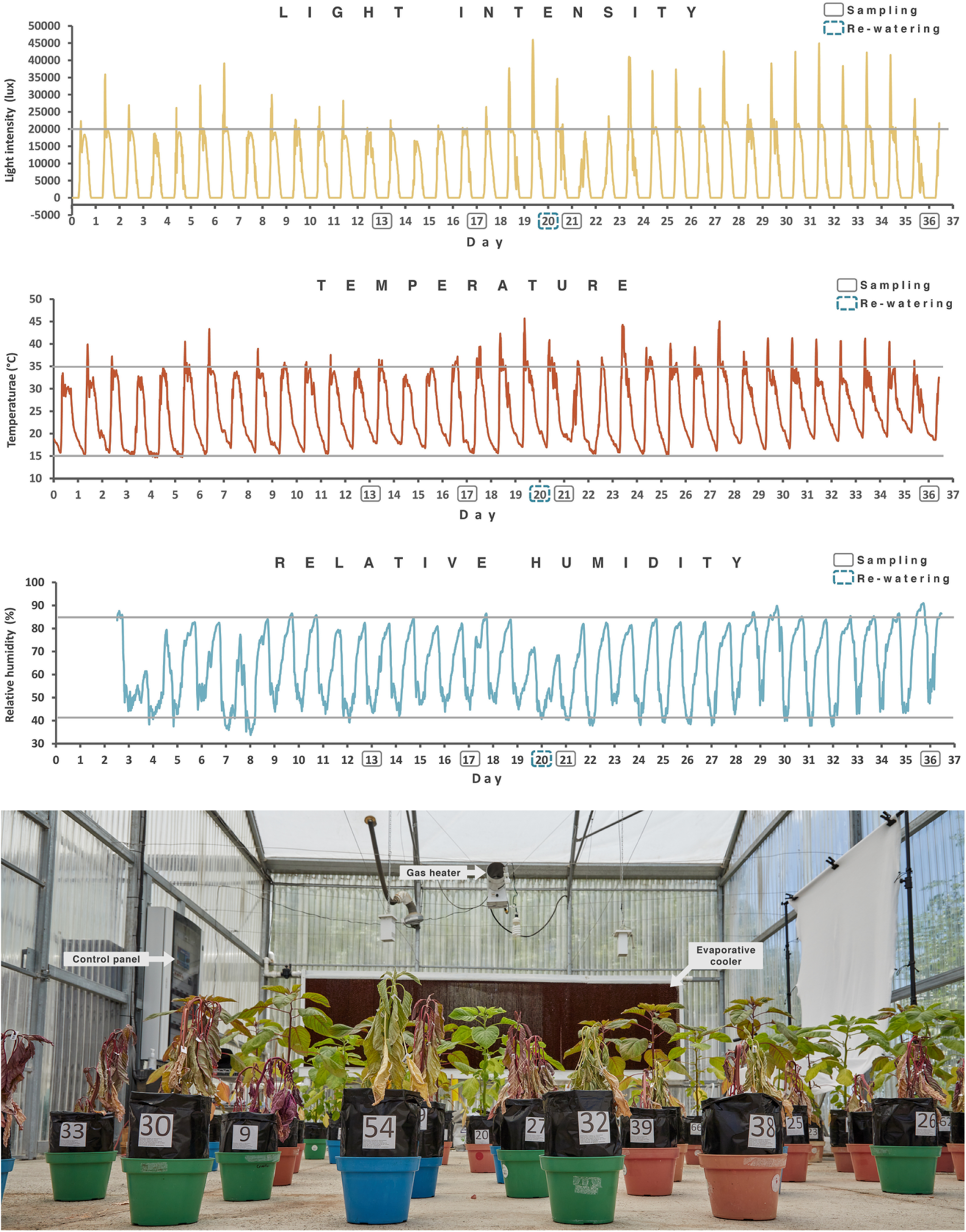
Growth conditions were controlled during the experiment. *Amaranthus cruentus* plants were germinated and grown for 20 days in growth chambers, then transplanted to 2 kg substrate and transferred to an acclimatized greenhouse equipped with a gas heater and evaporative cooler. Conditions during summer reached up to 45 °C at peak hours, but overall temperature (15–35 °C), light (≤45,000 lux), and relative humidity (40–85%) remained within controlled and homogeneous ranges.

**SUPPLEMENTARY FIGURE 2.**
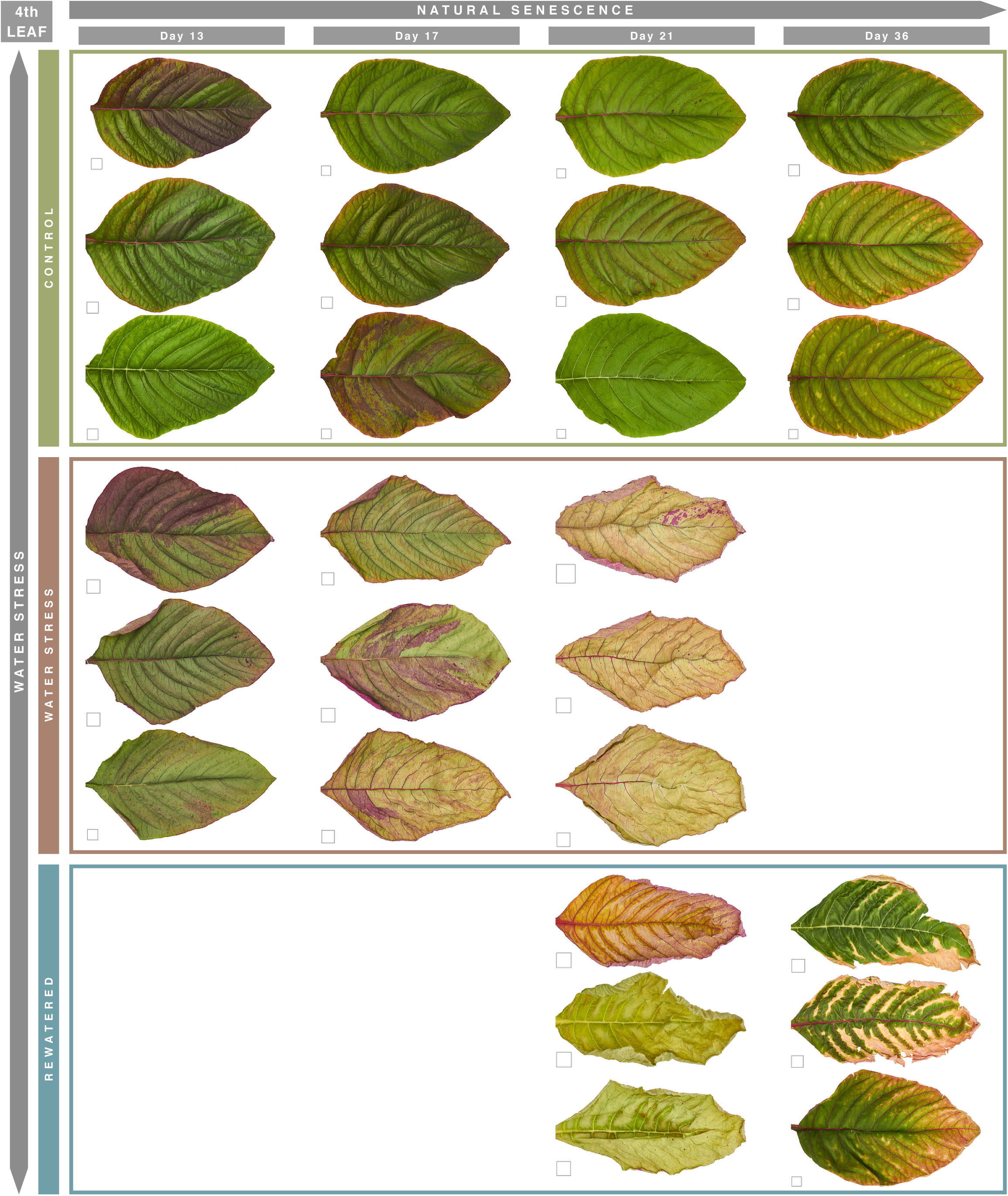
Response of 4th leaves to water stress. Control leaves remained young on Day 13 and began senescing by Day 36. Stressed leaves lost turgor and became almost completely yellow by Day 21, although they were not brittle. Rewatered leaves experienced similar stress but regained turgidity in major veins and adjacent lamina within 24 h (Day 21), and partially regreened by Day 36. Square = 1 x 1 cm.

**SUPPLEMENTARY FIGURE 3.**
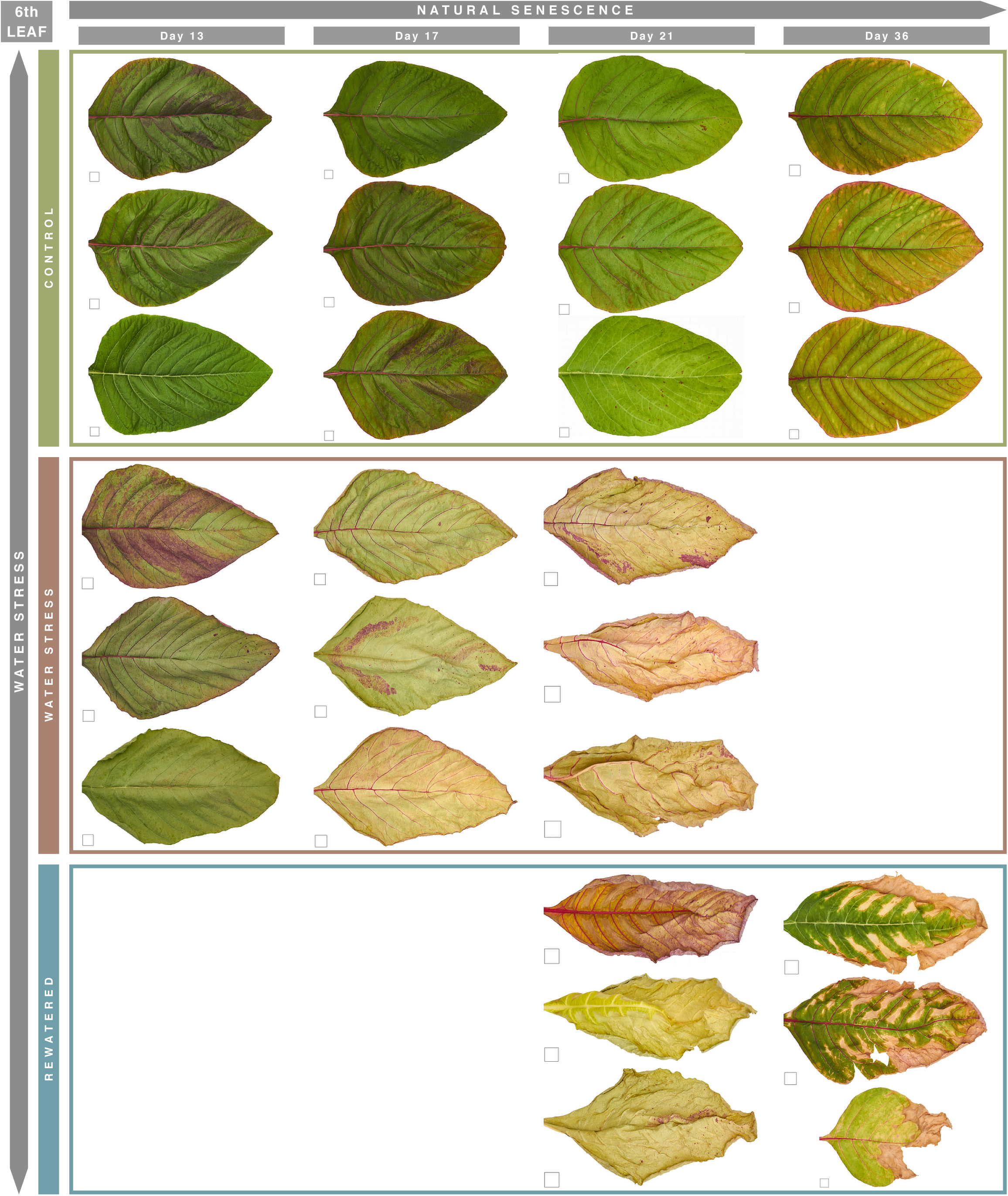
Response of 6th leaves to water stress. Control leaves were young on Day 13 and showed signs of senescence by Day 36. Stressed leaves lost turgor and became yellow and slightly brittle by Day 21. Rewatered leaves regained turgidity gradually after irrigation (Day 21) and partially regreened by Day 36, although less extensively than the 4th leaves. Square = 1 x 1 cm.

**SUPPLEMENTARY FIGURE 4.**
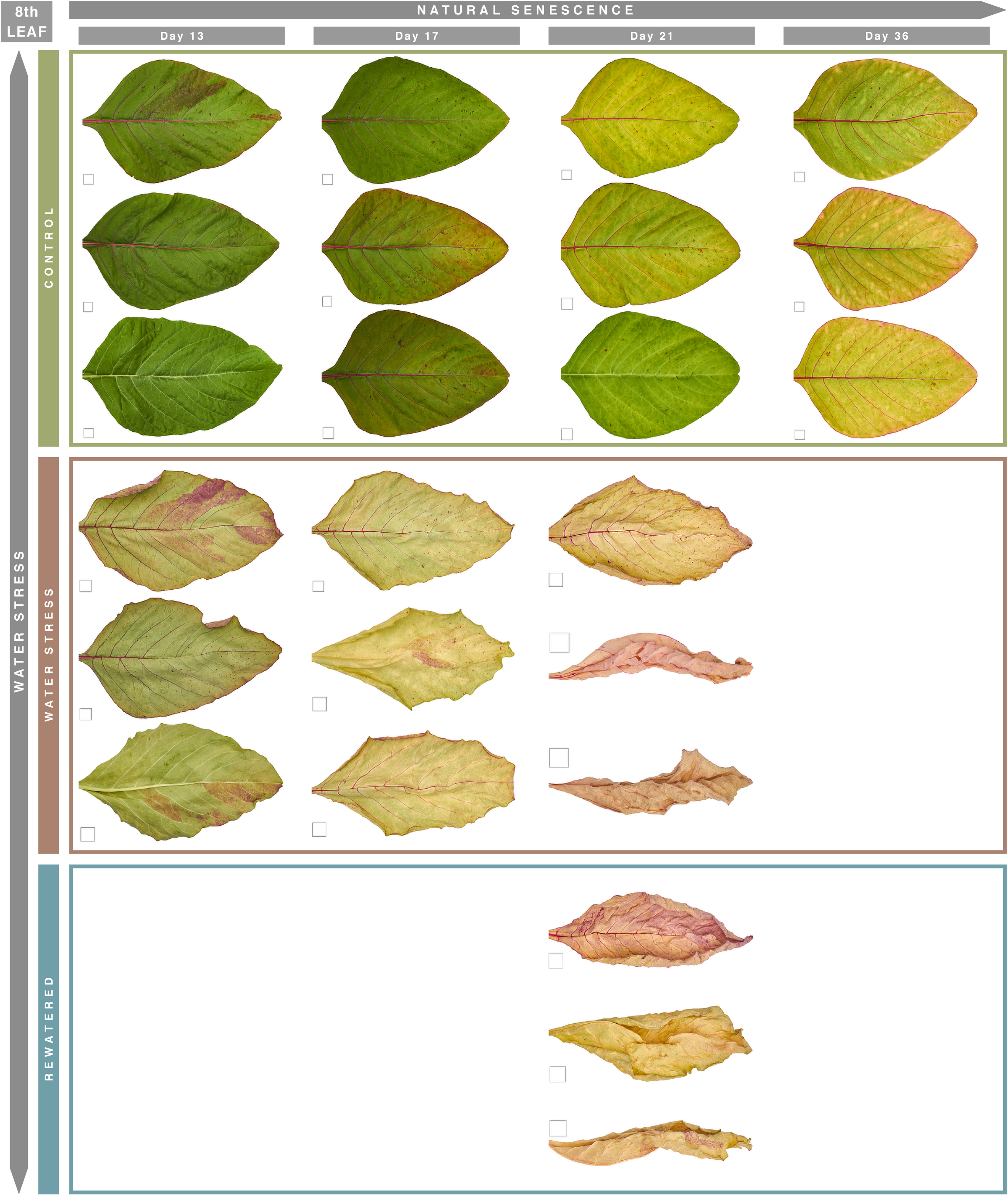
Response of 8th leaves to water stress. Control leaves began senescing by Day 13 and reached advanced senescence by Day 36. Stressed leaves lost turgor and became yellow and brittle by Day 21. Rewatered leaves did not recover and remained similar to stressed leaves. Square = 1 x 1 cm.

**SUPPLEMENTARY FIGURE 5.**
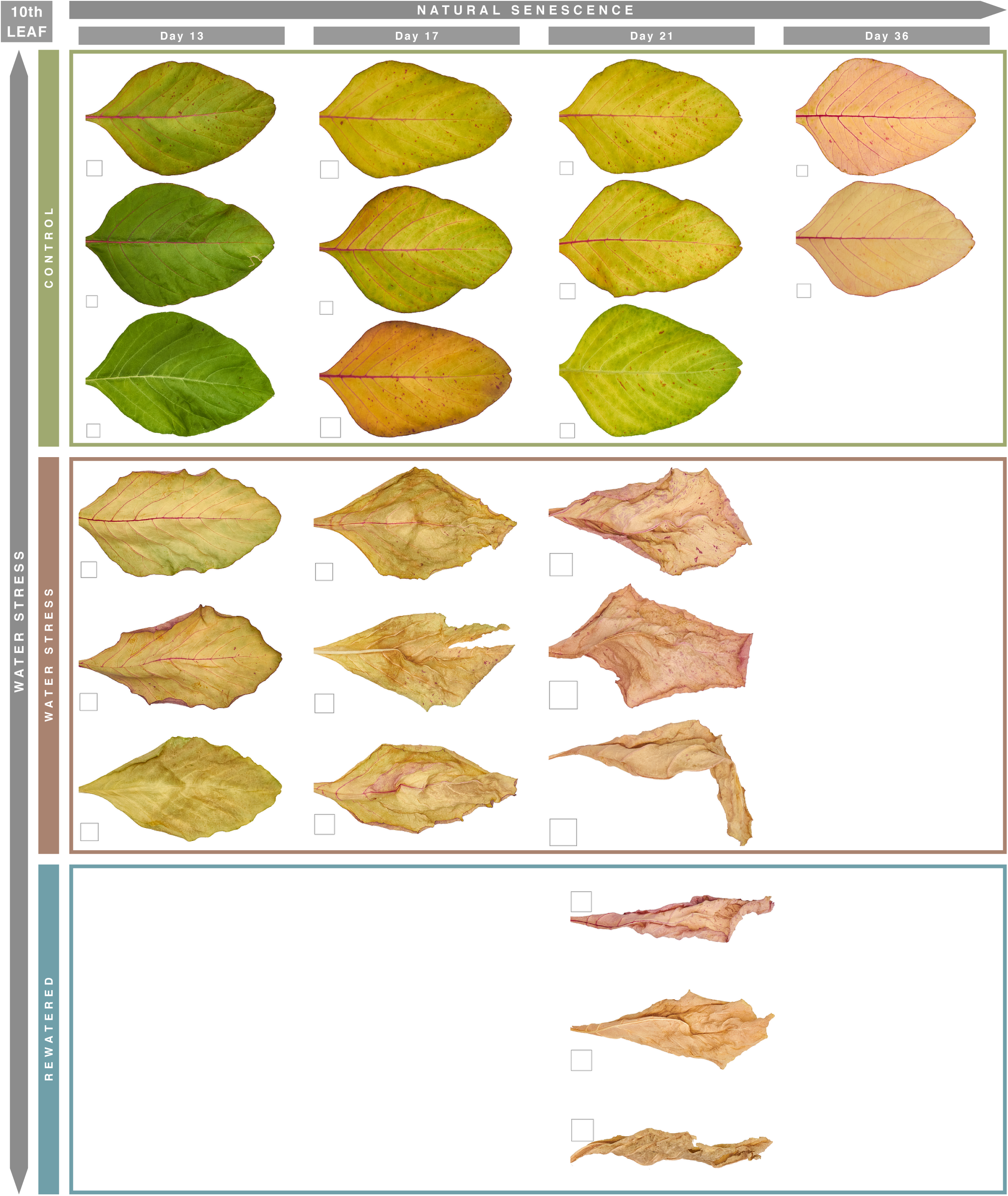
Response of 10th leaves to water stress. Control leaves were already senescing on Day 13 and reached complete senescence by Day 36, just before abscising. Stressed leaves lost turgor and became yellow and brittle by Day 21. Rewatered leaves did not recover and remained identical to stressed leaves. Naturally abscised leaves were not collected. Square = 1 x 1 cm.

**SUPPLEMENTARY FIGURE 6.**
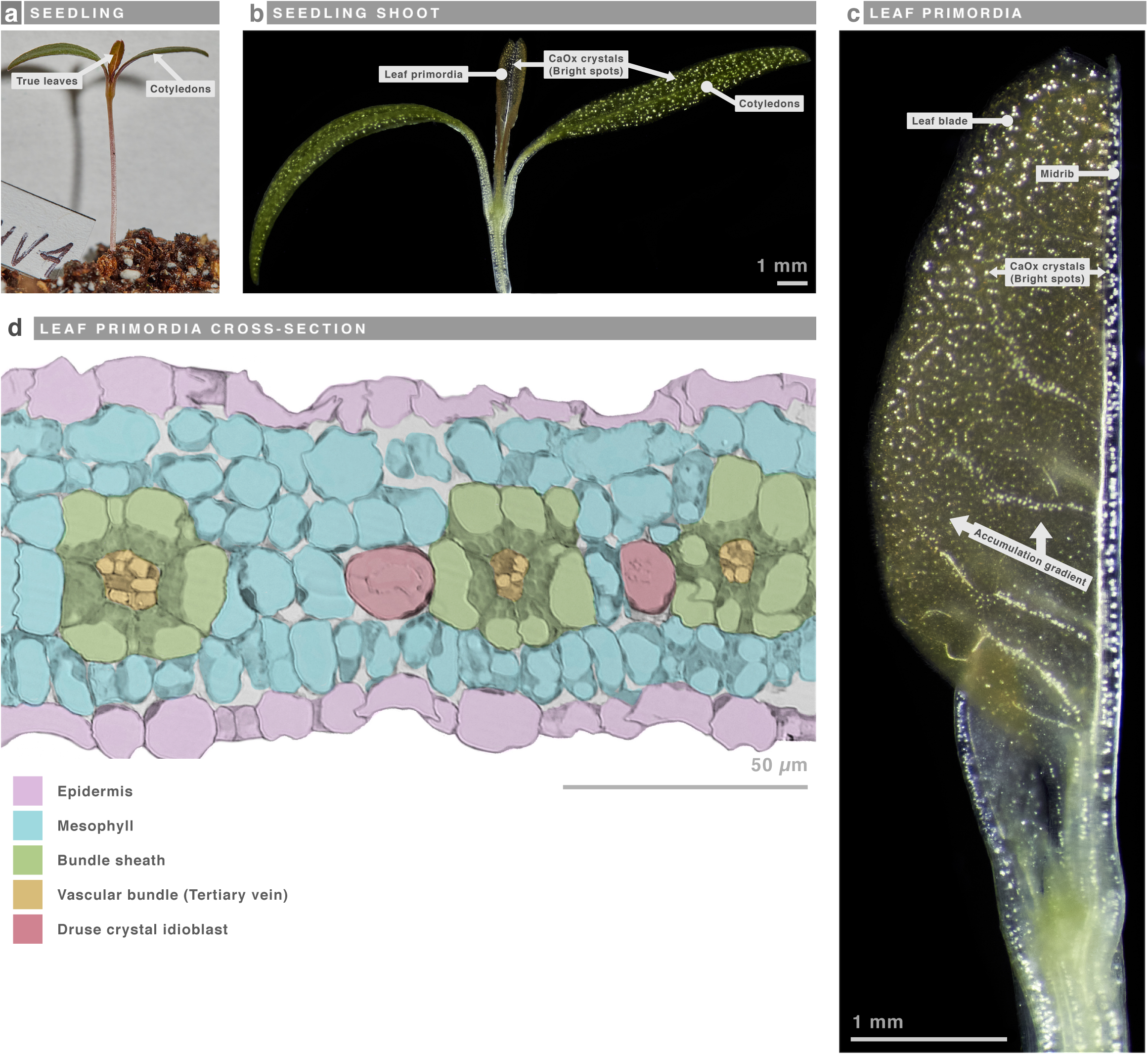
Calcium oxalate crystals appear early in leaf development. (a) Ten-day-old amaranth seedling. (b–c) Polarized-light micrographs of the shoot and leaf primordia. (d) DIC cross-section of a primordium (false-colored). CaOx crystals were present in all seedling organs, including epicotyl, cotyledons, and leaf primordia. In primordia, crystals appeared progressively from the midrib toward the margins, with one druse idioblast occurring between each vascular bundle.

**SUPPLEMENTARY FIGURE 7.**
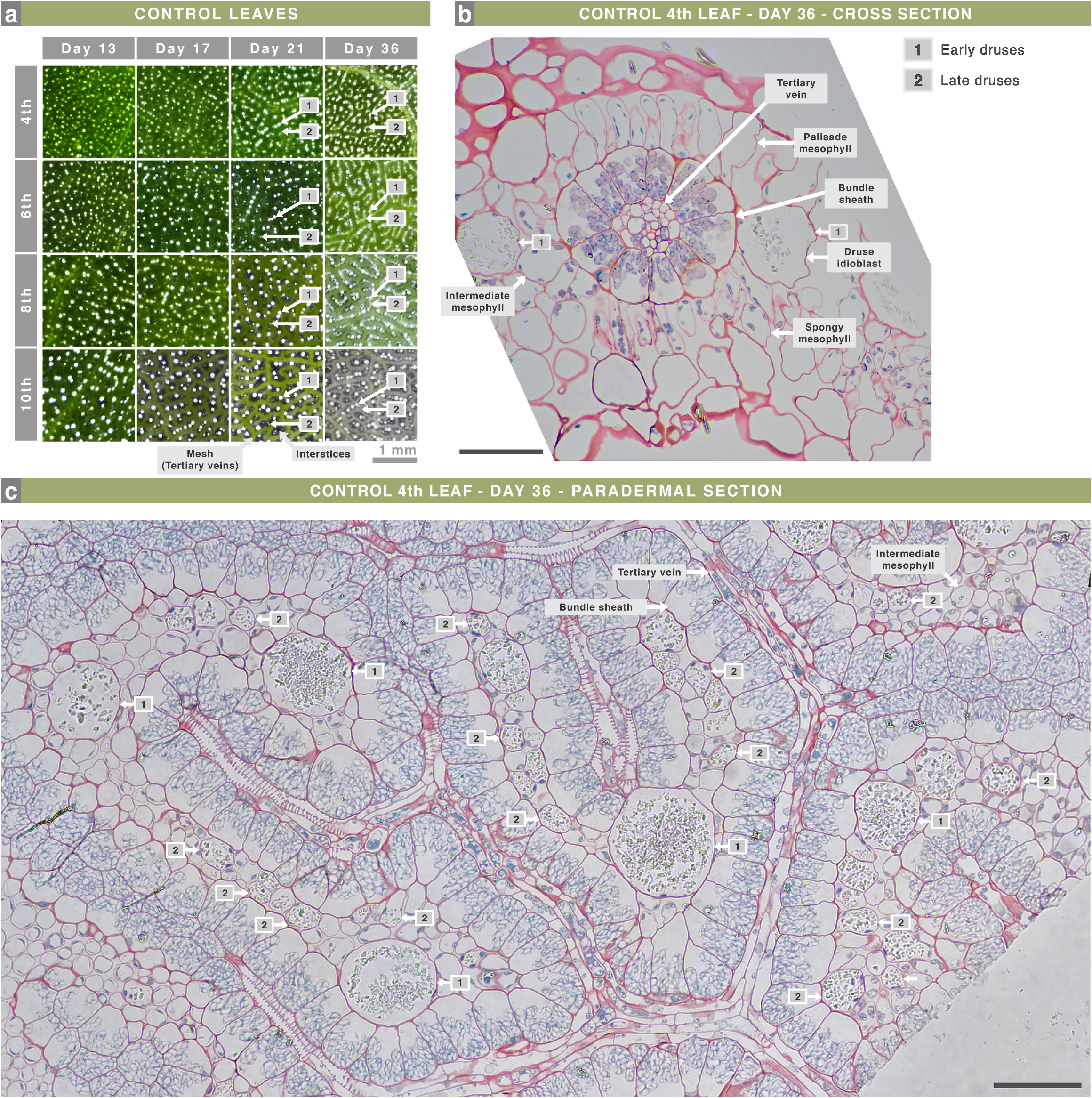
Intermediate mesophyll cells become crystal idioblasts during aging. (a) Polarized-light micrographs of fixed control leaves. (b) DIC cross-section of the 4th control leaf at Day 36. (c) DIC paradermal section of the same leaf. Tertiary veins form a mesh whose interstices are filled by intermediate mesophyll and druse idioblasts. By Day 13, large homogeneous druses (≈50 µm; “early druses”) predominated. During aging, smaller druses (10–30 µm; “late druses”) emerged between large druses, forming continuous linear arrangements. Both druse types occurred exclusively among intermediate mesophyll, which occupies the middle half of the leaf thickness. Leaf primordia contained only early druses, whereas late druses originated from intermediate mesophyll cells that dedifferentiated into crystal idioblasts as the leaf aged.

**SUPPLEMENTARY FIGURE 8.**
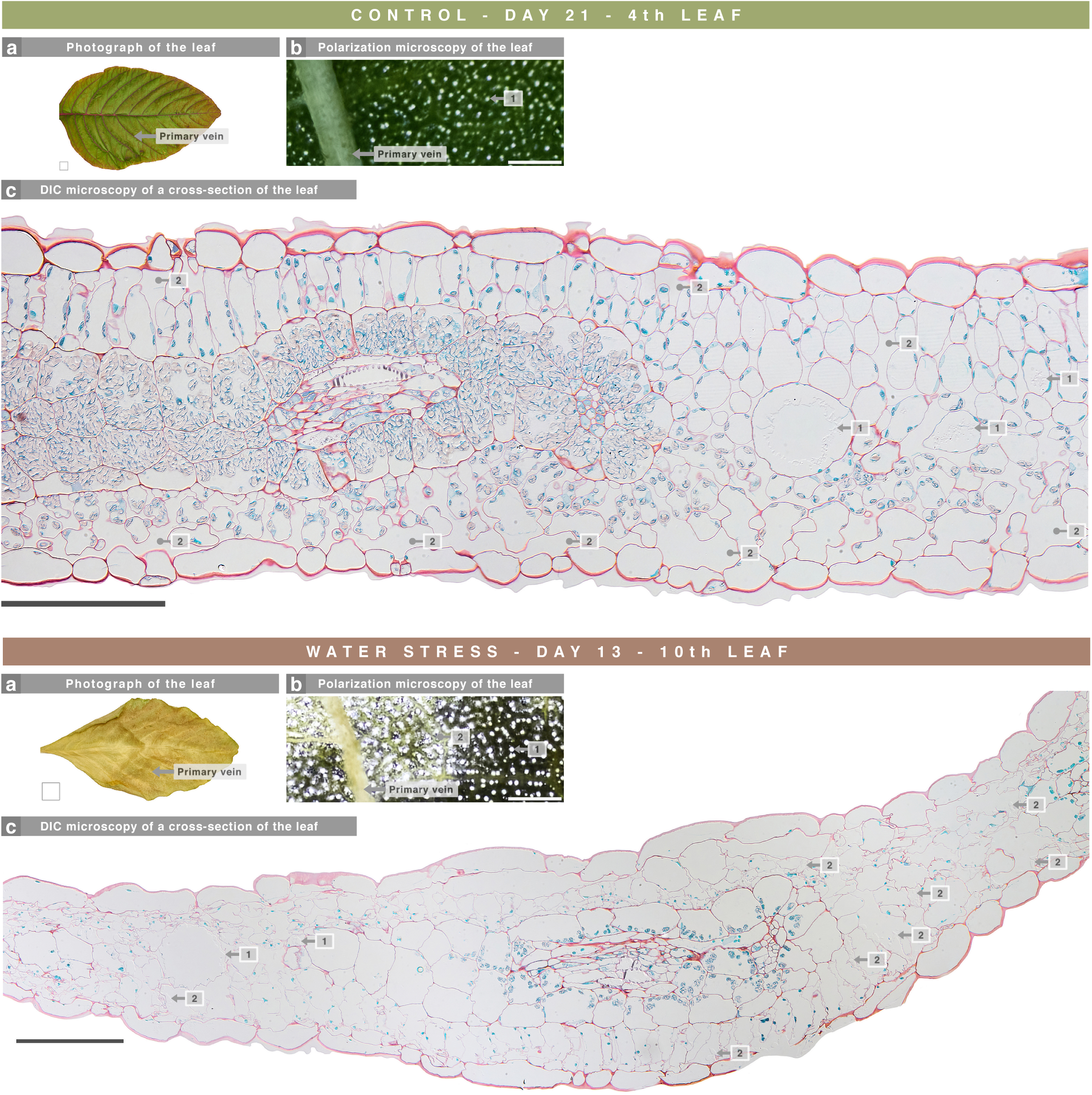
Water stress triggered extracellular accumulation of CaOx crystals. Photograph (a), polarized-light micrographs (b), and DIC sections (c) of the 4th control leaf (Day 21) and the 10th stressed leaf (Day 13). CaOx crystals are lost during sectioning, but their former locations appear as voids in resin under DIC (1). Water stress induced an abnormal and highly heterogeneous accumulation of crystals, mainly around primary veins in the 10th leaf, occurring both intracellularly in idioblasts (1) and, unusually, in extracellular spaces (2).

